# The helper NLR immune protein NRC3 mediates the hypersensitive cell death caused by the cell-surface receptor Cf-4

**DOI:** 10.1101/2021.09.28.461843

**Authors:** Jiorgos Kourelis, Mauricio P. Contreras, Adeline Harant, Hiroaki Adachi, Lida Derevnina, Chih-Hang Wu, Sophien Kamoun

## Abstract

Cell surface pattern recognition receptors (PRRs) activate immune responses that can include the hypersensitive cell death. However, the pathways that link PRRs to the cell death response are poorly understood. Here, we show that the cell surface receptor-like protein Cf-4 requires the intracellular nucleotide-binding domain leucine-rich repeat containing receptor (NLR) NRC3 to trigger a confluent cell death response upon detection of the fungal effector Avr4 in leaves of *Nicotiana benthamiana*. This NRC3 activity requires an intact N-terminal MADA motif, a conserved signature of coiled-coil (CC)-type plant NLRs that is required for resistosome-mediated immune responses. A chimeric protein with the N-terminal α1 helix of Arabidopsis ZAR1 swapped into NRC3 retains the capacity to mediate Cf-4 hypersensitive cell death. Pathogen effectors acting as suppressors of NRC3 can suppress Cf-4-triggered hypersensitive cell-death. Our findings link the NLR resistosome model to the hypersensitive cell death caused by a cell surface PRR.

## INTRODUCTION

Recognition of pathogens by the plant innate immune system predominantly relies on cell surface immune receptors and intracellular NOD-like/nucleotide-binding leucine-rich repeat receptors (NLRs). The activation of these receptors upon recognition of their cognate ligands triggers specific responses which ultimately result in activation of downstream immune pathways and disease resistance. The ligand sensing mechanisms are generally well-known, occurring either by direct binding of a pathogen-derived or pathogen-induced ligand to the receptor, or by indirect recognition requiring additional host proteins (Kourelis and van der Hoorn, 2018). Once cell-surface receptors and intracellular NLRs are activated, both classes of receptors induce common signaling pathways, such as Ca^2+^ influx, ROS production and MAPK activation. However, the molecular mechanisms by which receptor activation results in common downstream immune signaling are not as well understood. The extent to which the downstream signaling of cell surface immune receptors and intracellular NLRs differs and converges is one of the main open questions in the field of plant immunity (Lu and Tsuda, 2021).

The plant NLR family is defined by a central NB-ARC (nucleotide-binding domain shared with APAF-1, various R-proteins and CED-4) domain and at least one other domain (Kourelis *et al*., 2021). Plant NLRs can be subdivided in four main subclades characterized by distinct N-terminal domains. TIR-NLRs contain a catalytically active N-terminal Toll/interleukin-1 receptor (TIR) domain, while CC_R_-NLRs, CC_G10_-NLRs, and CC-NLRs contain distinct RPW8-type, G10-type, or Rx-type coiled coil (CC) domains, respectively. Much progress has been made in recent years in understanding the molecular mechanisms of recognition and signaling of plant NLRs. In general, NLR activation upon ligand recognition results in an exchange of ADP to ATP in the NB-ARC domain and the formation of an oligomeric complex, termed a “resistosome”. The Arabidopsis CC-NLR ZAR1 forms a pentameric resistosome structure in which the N-terminal α1 helix extends out to form a funnel which acts as a Ca^2+^-permeable channel (Wang, Hu, *et al*., 2019; Wang, Wang, *et al*., 2019; Bi *et al*., 2021). Similarly, the CC_R_-NLRs NRG1.1 and ADR1 generate Ca^2+^-permeable channels upon activation (Jacob *et al*., 2021). The TIR-NLRs RPP1 and Roq1 form tetrameric resistosome complexes upon activation (Ma *et al*., 2020; Martin *et al*., 2020). The oligomerization of the TIR domain results in NADase activity and the release of a plant-specific variant cyclic ADP-ribose (v-cADPR) (Horsefield *et al*., 2019; Wan *et al*., 2019). A subset of NLRs, known as functional singletons, combine pathogen detection and immune signaling activities into a single protein (Adachi, Derevnina, *et al*., 2019). In contrast, many NLRs have functionally specialized over evolutionary time in “sensor” or “helper” NLRs dedicated to pathogen recognition or immune signaling, respectively (Adachi, Derevnina, *et al*., 2019). Sensor and helper NLRs function together and can form receptor networks with a complex architecture (Wu *et al*., 2017; Wu *et al*., 2018).

Cell surface receptors are typically divided in two categories: the receptor-like kinases (RKs, also known as RLKs) and the receptor-like proteins (RPs, also known as RLPs). RKs contain a C-terminal intracellular kinase domain for downstream signaling, while RPs contain a small cytoplasmic tail and lack a C-terminal kinase domain. Both RKs and RPs have a single-pass transmembrane domain and a variety of extracellular ligand-binding domains (Zipfel, 2014). Upon ligand perception, RKs typically hetero-oligomerize with other RKs which act as coreceptors. For example, binding of chitin by the Arabidopsis LysM-RK LYK5 induces hetero-oligomerization with the LysM-RK co-receptor CERK1 (Cao *et al*., 2014). Furthermore, recognition of the flg22 peptide of bacterial flagellin by the leucine-rich repeat (LRR)-RK FLS2 induces hetero-oligomerization with the LRR-RK co-receptor SERK3/BAK1 (Sun *et al*., 2013). Similarly, most LRR-RPs constitutively interact with the LRR-RK SOBIR1, and activation upon ligand binding induces the hetero-oligomerization with the LRR-RLK co-receptor SERK3/BAK1 (Liebrand *et al*., 2013; Zhang *et al*., 2014; Albert *et al*., 2015; Bi *et al*., 2016; Postma *et al*., 2016; van der Burgh *et al*., 2019; Huang *et al*., 2021). These activated complexes in turn phosphorylate specific downstream receptorlike cytoplasmic kinases (RLCKs) which ultimately relay the immune response (Lu and Tsuda, 2021; DeFalco and Zipfel, 2021).

While the signaling cascades downstream of cell surface immune receptors and NLRs were thought to be distinct (Jones and Dangl, 2006), it is now becoming clear that in fact these signaling modules intertwine at different points (Ngou *et al*., 2021; Yuan *et al*., 2021; Pruitt *et al*., 2021). Cell surface immune signaling was recently shown to be required for the hypersensitive cell death immune response triggered by a variety of NLRs (Yuan *et al*., 2021; Ngou *et al*., 2021). Specifically, in the absence of cell surface immune signaling, the activation of the TIR-NLR pair RRS1/RPS4 by the effector AvrRps4 does not result in hypersensitive cell death (Ngou *et al*., 2021). Moreover, activation of the CC_G10_-NLRs RPS2 and RPS5 by AvrRpt2 (Yuan *et al*., 2021; Ngou *et al*., 2021) and AvrPphB (Ngou *et al*., 2021), respectively, or the CC-NLR RPM1 by AvrRpm1 (Ngou *et al*., 2021) results in diminished hypersensitive cell death in the absence of cell surface immune signaling. The emerging model is that cell surface immune signaling and intracellular NLR signaling mutually potentiate each other by enhancing transcription of the core signaling components of these pathways (Ngou *et al*., 2021; Yuan *et al*., 2021). However, the mechanistic links between cell surface immune receptors and NLRs are not yet known (Bjornson and Zipfel, 2021). Additionally, there is a tendency in the field to view cell surface and NLR receptors through a unified lens despite their huge structural and functional diversity (Kourelis *et al*., 2021; Zipfel, 2014). The degree to which distinct classes of cell surface and intracellular receptors engage in signaling networks remains unclear.

In addition to their potentiation of cell surface immune signaling, NLRs can also be genetically involved downstream of cell surface immune receptor activation. In Arabidopsis, the EDS1/PAD4/ADR1 and, to a lesser extent, EDS1/SAG101/NRG1 modules are genetically required for a subset of the immune responses triggered by LRR-RPs (Pruitt *et al*., 2021; Tian *et al*., 2021) and LRR-RKs (Tian *et al*., 2021). However, the hypersensitive cell death triggered by a subset of LRR-RPs is not affected in these mutants (Pruitt *et al*., 2021). Since the EDS1/PAD4/ADR1 and EDS1/SAG101/NRG1 modules are typically associated with hypersensitive cell death signaling, this raises the question of what the molecular mechanism of this interaction is. Indeed, the extent to which the subset of LRR-RPs that trigger hypersensitive cell death require NLRs warrants further investigation. Previously, the NLR protein NRC1 was implicated in the cell death mediated by the LRR-RLPs Cf-4 (Gabriëls *et al*., 2006), LeEIX2 (Gabriëls *et al*., 2007), and Ve1 (Fradin *et al*., 2009). However, these studies were based on virus-induced gene silencing and, therefore, off-target effects on other NLR genes cannot be ruled out. Indeed, the silencing experiments performed in *Nicotiana benthamiana* were based on a fragment sequence corresponding to the LRR domain of tomato NRC1 and predated the sequencing of the *N. benthamiana* genome (Wu *et al*., 2016). The sequencing of the *N. benthamiana* genome revealed that *N. benthamiana* does not encode a functional ortholog of the tomato NRC1 gene (Bombarely *et al*., 2012; Naim *et al*., 2012; Nakasugi *et al*., 2014). Instead, the *N. benthamiana* genome contains homologs of NRC1, namely NRC2, NRC3, and NRC4 (Wu *et al*., 2017). Therefore, the precise contribution of NLRs to the hypersensitive response mediated by Cf-4 type receptors has remained unclear, due also to the weak phenotypes observed using gene silencing constructs targeting the various *N. benthamiana* NRCs (Wu *et al*., 2016; Brendolise *et al*., 2017).

The NRC superclade is a subclade of CC-NLRs that has massively expanded in some asterid plants, including the Solanaceae (Wu *et al*., 2017). The NRC superclade consists of *R* gene encoded sensor NLRs which confer resistance to diverse pathogens, and the NRC (NLR required for cell death) helper CC-NLRs, which mediate immune signaling and the hypersensitive cell death response downstream of effector recognition by the sensor NLRs. NRCs form redundant nodes in complex receptor networks, with different NRCs exhibiting different specificities for their sensor NLRs. Much like ZAR1, the N-terminal α1 helix of NRCs is defined by a molecular signature called the “MADA motif’ (Adachi, Contreras, *et al*., 2019). NRCs and ZAR1 are therefore classified as MADA-CC-NLRs, a class that includes ~20% of all CC-NLRs in angiosperms (Adachi, Contreras, *et al*., 2019; Kourelis *et al*., 2021). In solanaceous species, the NRC network comprises up to half of the NLRome and can be counteracted by pathogen effectors. Recently, Derevnina *et al*., (2021) showed that effector proteins from the cyst nematode *Globodera rostochiensis* and the oomycete *Phytophthora infestans* specifically suppress the activity of NRC2 and NRC3, independently of their sensor NLR partners.

In this study, we took advantage of various combinations of CRISPR/Cas9-induced loss-of-function mutations in the *N. benthamiana* NRCs to revisit the connection between NRCs and cell surface receptor (Adachi, Contreras, *et al*., 2019; Wu *et al*., 2020; Witek *et al*., 2021). We hypothesized that an activated NRC resistosome is required downstream of LRR-RP activation in order to produce hypersensitive cell death. To challenge this hypothesis, we used the LRR-RP Cf4 (Thomas *et al*., 1997), which upon recognition of the fungal pathogen *Cladosporium fulvum* (Syn. *Passalora fulva*) apoplastic effector Avr4 triggers hypersensitive cell death (Joosten *et al*., 1994). Using a combination of reverse genetics and complementation, we determined that Cf-4/Avr4-triggered cell death is largely mediated by NRC3, and not NRC1. Using defined mutations in the MADA motif and by generating swaps with the Arabidopsis ZAR1 α1 helix, we show that NRC resistosome function is likely downstream of Cf-4 activation by Avr4. Finally, we show that divergent pathogen-derived suppressors of NRC activity suppress Cf-4/Avr4-triggered cell death. We conclude that Cf-4/Avr4-triggered hypersensitive cell death is mediated by the MADA-CC-NLR NRC3, likely requiring the formation of a Ca^2+^-permeable pore. Divergent pathogens target this node, thereby suppressing both intracellular and cell surface triggered immune responses. NRC3 is conserved in the Solanaceae and likely mediates hypersensitive cell death triggered by a variety of leucine-rich repeat receptor-like proteins. We propose that NRC3 is a core node that connects cell surface and intracellular immune networks.

## RESULTS

### Cf-4/Avr4-triggered hypersensitive cell death is compromised in the *N. benthamiana nrc23* CRISPR mutant line

We generated multiple mutant lines of *N. benthamiana* that carry CRISPR/Cas9-induced premature termination mutations in various combinations of the *NRC2* (*NRC2a* and *NRC2b*), *NRC3* and *NRC4* (*NRC4a* and *NRC4b*) genes. These lines enabled us to revisit the contribution of NRC helper NLRs to the hypersensitive cell death caused by the receptor-like protein Cf-4. To do this, we quantified the cell death response following transient expression of Cf-4/Avr4 by agroinfiltration, or as controls the receptor/effector pairs Pto/AvrPto (mediated by Prf; NRC2/3-dependent), Rpi-blb2/AVRblb2 (NRC4-dependent), or R3a/Avr3a (NRC-independent) (**Figure 1, Figure S1**) (Wu *et al*., 2017). Cf-4/Avr4-triggered cell-death was significantly reduced in both the *nrc2/3*, and *nrc2/3/4* CRISPR mutant lines as compared to wild-type (**Figure 1A**). Cf-4 protein accumulation was not affected in the *nrc2/3/4* CRISPR lines (**Figure 1B**), indicating that the reduction in cell death is not correlated with Cf-4 protein level. While Pto/AvrPto-triggered cell death is completely abolished in the *nrc2/3* and *nrc2/3/4* CRISPR lines (**Figure 1A, C**, **Figure S1B**), Cf-4/Avr4 expression resulted in a specific chlorosis in the *nrc2/3* and *nrc2/3/4* lines which did not develop into confluent cell death (**Figure 1C**, **Figure S1B**). These results indicate that the Cf-4/Avr4-triggered hypersensitive cell death requires NRC2/3.

**Figure 1:**
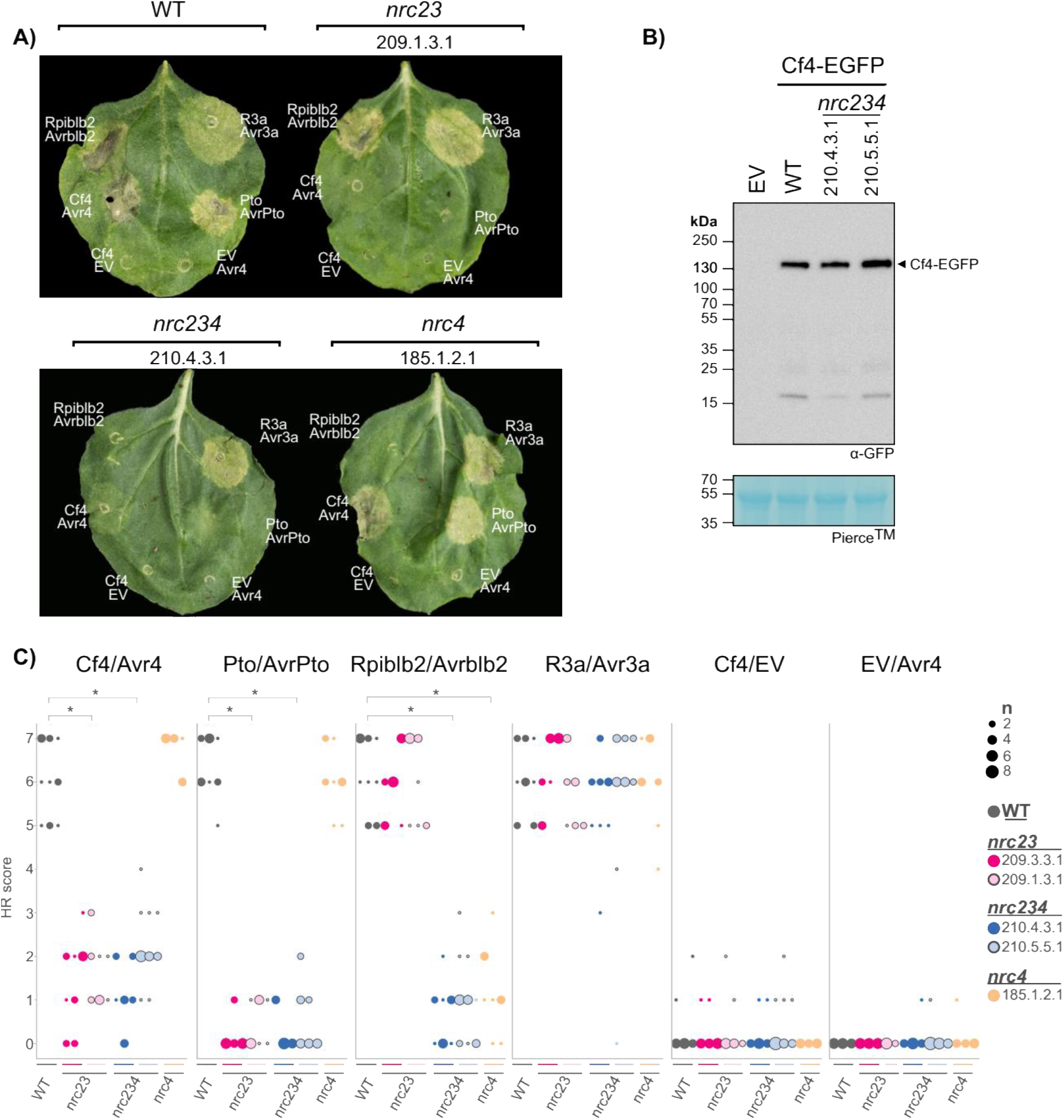
Cf-4/Avr4-triggered hypersensitive cell death requires NRC2/3. **A**) Cf-4/Avr4-triggered hypersensitive cell death is lost in the *nrc2/3* and *nrc2/3/4* knock-out lines, but not in the *nrc4* line. Representative *N. benthamiana* leaves infiltrated with appropriate constructs photographed 7 – 10 days after infiltration. The NRC CRISPR lines, *nrc2/3*-209.1.3.1, *nrc2/3/4*-210.4.3.1, and *nrc4*-185.1.2.1, are labelled above the leaf and the receptor/effector pair tested, Cf-4/Avr4, Prf (Pto/AvrPto), Rpi-blb2/AVRblb2 or R3a/Avr3a, are labelled on the leaf image. Cf-4/EV and EV/Avr4 were also included as negative controls. A representative leaf of the independent *nrc2/3*-209.3.3.1, and *nrc2/3/4*-210.5.5.1 CRISPR line are shown in **Figure S1**. **B**) Cf-4 protein accumulation is not affected in the *nrc2/3/4* lines. For the immunoblot analysis, total protein was extracted 5 days after transient expression of Cf-4-EGFP by agroinfiltration in wild-type, *nrc2/3/4*-210.4.3.1 and *nrc2/3/4*-210.5.5.1 *N. benthamiana* leaves. Cf-4-EGFP accumulation was detected using anti-GFP antibody. **C**) Quantification of hypersensitive cell death. Cell death was scored based on a 0 - 7 scale (**Figure S1**) between 7 – 10 days post infiltration. The results are presented as a dot plot, where the size of each dot is proportional to the count of the number of samples with the same score within each biological replicate. The experiment was independently repeated three times. The columns correspond to the different biological replicates. Significant differences between the conditions are indicated with an asterisk (*). Details of statistical analysis are presented in **Figure S1**.

### NRC3 mediates Cf-4/Avr4-triggered hypersensitive cell death in *N. benthamiana*

Next, we determined the individual contribution of NRC2 and NRC3 to Cf-4/Avr4-triggered cell death. To do this, we transiently complemented the *nrc2/3/4* CRISPR lines with either NRC2 or NRC3 in addition to expressing Cf-4/Avr4 (**Figure 2A**). As a control, we used Pto/AvrPto, which can utilize either NRC2 or NRC3 for Prf-mediated hypersensitive cell death signalling in *N. benthamiana* (Wu *et al*., 2016). In addition to NRC2 and NRC3, we used NRC4 which is not expected to complement the Cf-4/Avr4 or Pto/AvrPto-triggered cell death in the *N. benthamiana nrc2/3/4* lines. Although it was previously reported that NRC1 is required for Cf-4/Avr4 hypersensitive cell death (Gabriëls *et al*., 2006), *N. benthamiana* and other species in the *Nicotiana* genus lack *NRC1* (**Figure S2**). Thus, we decided to use tomato NRC1 (SlNRC1) to test whether *Sl*NRC1 can also complement Cf-4/Avr4-triggered cell death in *N. benthamiana*. In the *nrc2/3/4* lines, Cf-4/Avr4 and Pto/AvrPto-triggered cell death responses were complemented by NRC3 compared to the empty vector (EV) or NRC4 control (**Figure 2B, C**, **Figure S3A** for statistical analysis). NRC2 complemented Pto/AvrPto cell death in the *nrc2/3/4*, but only weakly complemented Cf-4/Avr4-triggered cell death (**Figure 2B, C**). Finally, *Sl*NRC1 weakly complemented Pto/AvrPto cell death, but not Cf-4/Avr4 cell death **Figure 2B, C**, **Figure S3A** for statistical analysis. Whereas NRC1 is pseudogenized in the *Nicotiana* genus with only small fragments corresponding to the LRR being found in the genome assemblies, NRC2 and NRC3 are conserved in all Solanaceous species investigated (**Figure S2**). We conclude that NRC3 is the main NRC helper node contributing to Cf-4/Avr4-triggered hypersensitive cell death in *N. benthamiana*.

**Figure 2:**
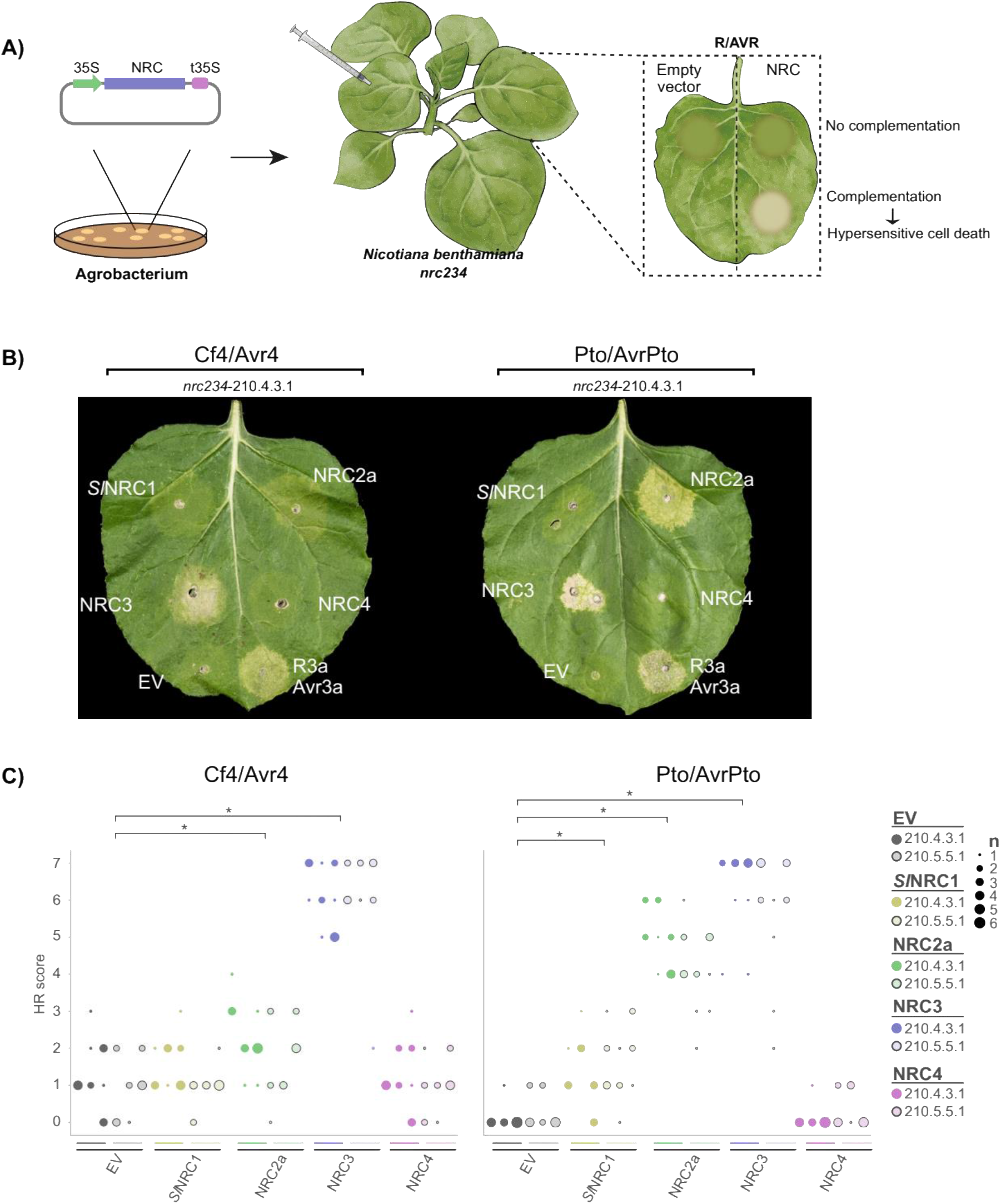
NRC3 complements Cf-4/Avr4-triggered hypersensitive cell death in the *nrc2/3/4* knock-out lines. **A**) A schematic representation of the cell death complementation assay. NRCs and empty vector (EV) were transformed into *A. tumefaciens*, and transiently co-expressed *in N. benthamiana* plants with either Cf-4/Avr4 or Pto/AvrPto. Hypersensitive cell death was scored based on a modified 0 - 7 scale between 7 – 10 days post infiltration (**Figure S1a**). **B**) Cf-4/Avr4-triggered hypersensitive cell death is complemented in the *nrc2/3/4* lines with NRC3 and to a lesser extent NRC2, but not *Sl*NRC1 or NRC4. Representative *N. benthamiana* leaves infiltrated with appropriate constructs were photographed 7 – 10 days after agroinfiltration. The receptor/effector pair tested, Cf-4/Avr4 and Prf (Pto/AvrPto), are labelled above the leaf of *nrc2/3/4*-210.4.3.1. The NRC used for complementation or EV control are labelled on the leaf image. A representative leaf of the independent *nrc2/3/4*-210.5.5.1 line is shown in **Figure S3**. **C**) Quantification of hypersensitive cell death. Cell death was scored based on a 0 - 7 scale between 7 – 10 days post infiltration. The results are presented as a dot plot, where the size of each dot is proportional to the count of the number of samples with the same score within each biological replicate. The experiment was independently repeated three times. The columns correspond to the different biological replicates. Significant differences between the conditions are indicated with an asterisk (*). Details of statistical analysis are presented in **Figure S3**.

### The NRC3 N-terminal MADA motif is required for Cf-4-triggered hypersensitive cell death

The NRC-helper clade is characterized by a N-terminal MADA-type α1 helix diagnostic of ZAR1-type CC-NLRs (Adachi, Contreras, *et al*., 2019). In order to determine whether the N-terminal MADA motif of NRC3 is required for the Cf-4/Avr4-triggered hypersensitive cell death, we took advantage of recently described cell death abolishing “MADA” mutations in this motif (Adachi, Contreras, *et al*., 2019). Like ZAR1 and NRC4, both NRC2 and NRC3 have an N-terminal MADA-type α1 helix (**Figure 3A**). Mutating leucine 17 to glutamic acid (L17E) abolishes NRC4-mediated cell death and immunity (Adachi, Contreras, *et al*., 2019). NRC2^L17E^ and NRC3^L21E^ correspond to the NRC4^L17E^ mutant (**Figure 3A, B**). To establish whether these mutations affect NRC2- or NRC3-mediated cell death, we conducted a complementation assay of Cf-4/Avr4 or Pto/AvrPto cell death in *N. benthamiana nrc2/3/4* lines with either wild type NRC2 or NRC3, or their respective MADA mutants (**Figure 3B**). The MADA mutation did not affect protein accumulation, as both NRC2, NRC3, and their respective MADA mutants accumulated to similar levels (**Figure 3C**). Unlike wild type NRC3, NRC3^L21E^ didn’t complement either Cf-4/Avr4 or Pto/AvrPto-triggered cell death in the *nrc2/3/4* lines (**Figure 3D, E**). As previously noted, NRC2 weakly complemented Cf-4/Avr4-triggered cell-death, but the NRC2^L17E^ failed to complement either this cell death or Pto/AvrPto-triggered cell death (**Figure 3D, E**, **Figure S4A** for the statistical analysis). We conclude that the N-terminal α1 helix of NRC3 is required for Cf-4/Avr4-triggered hypersensitive cell death.

**Figure 3:**
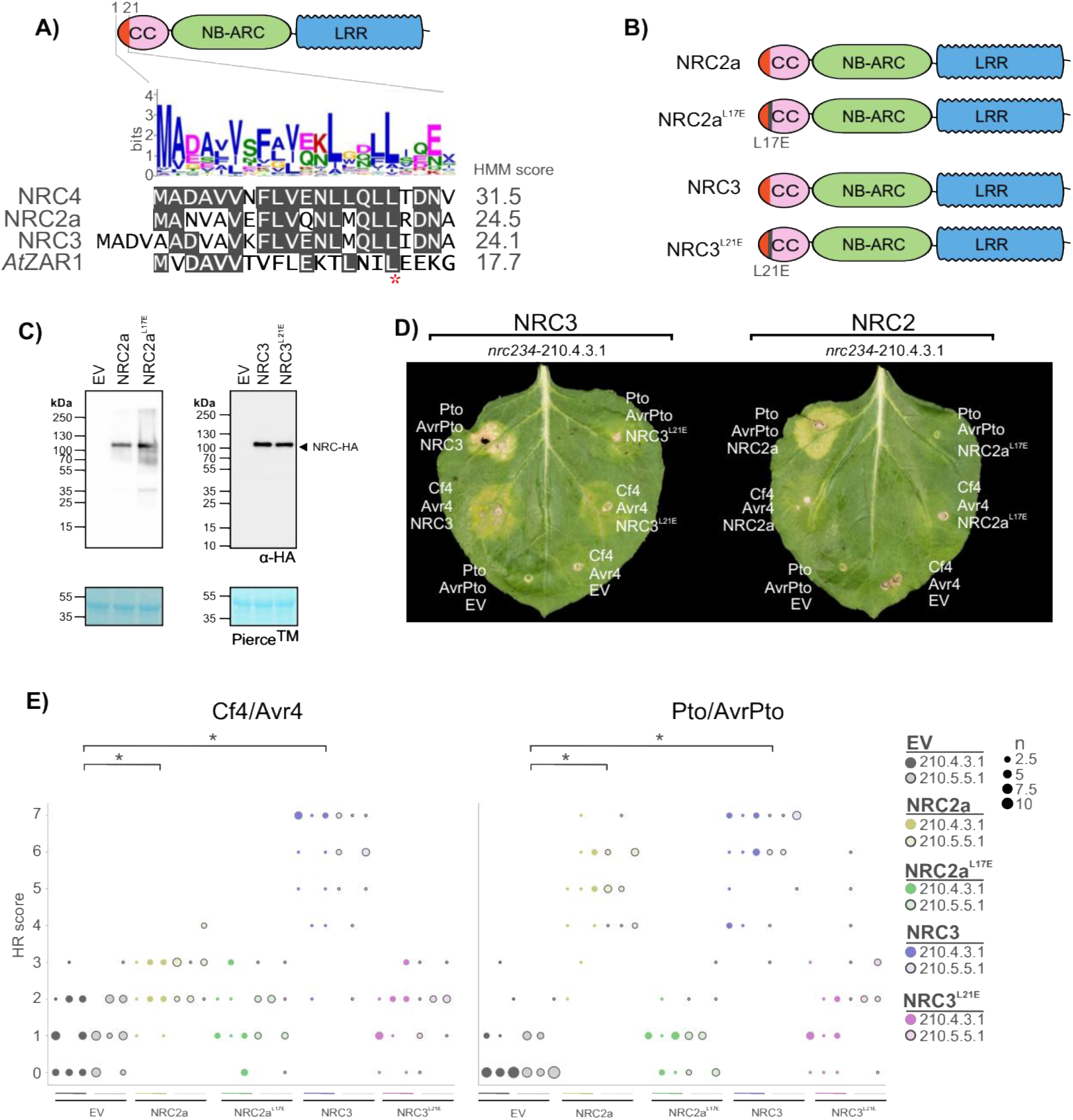
The N-terminal MADA motif of NRC3 is required for Cf-4/Avr4-triggered hypersensitive cell death. **A**) NRC3 has an N-terminal MADA motif. Alignment of NRC2, NRC3, NRC4 and ZAR1 N-terminal MADA motif with the consensus sequence pattern of the MADA motif and the HMM score for each sequence. **B**) Schematic representation of NRC2, NRC3, and the respective NRC2^L17E^ and NRC3^L21E^ MADA mutants. **C**) NRC2 and NRC3 MADA mutants accumulate to similar levels as wild type proteins. Immunoblot analysis of transiently expressed C-terminally 6xHA-tagged NRCs 5 days after agroinfiltration in wild-type *N. benthamiana* plants. **D**) NRC MADA mutants do not complement Cf-4/Avr4 or Pto/AvrPto-triggered hypersensitive cell death in the *N. benthamiana nrc2/3/4* lines. Representative *N. benthamiana* leaves infiltrated with appropriate constructs were photographed 7 – 10 days after infiltration. The NRCs tested, NRC2 and NRC3, are labelled above the leaf of *nrc2/3/4*-210.4.3.1. The receptor/effector pair tested, Cf-4/Avr4 and Prf (Pto/AvrPto), are labelled on the leaf image. A representative leaf of the independent *nrc2/3/4*-210.5.5.1 line is shown in **Figure S4**. **E**) Quantification of hypersensitive cell death. Cell death was scored based on a 0 - 7 scale between 7 – 10 days post infiltration. The results are presented as a dot plot, where the size of each dot is proportional to the count of the number of samples with the same score within each biological replicate. The experiment was independently repeated three times. The columns correspond to the different biological replicates. Significant differences between the conditions are indicated with an asterisk (*). Details of statistical analysis are presented in **Figure S4**.

### A chimeric protein fusion of the Arabidopsis ZAR1 α1 helix to NRC2/3 can complement loss of Cf-4-triggered hypersensitive cell death

In the activated ZAR1 resistosome, the N-terminal α1 helix of the CC domain forms a Ca^2+^-permeable membrane-permeable pore (Bi *et al*., 2021). In order to determine whether a similar mechanism also applies for Cf-4/Avr4-triggered hypersensitive cell death, we generated chimaeras of NRC2 or NRC3 (NRC2^ZAR1α1^ and NRC3^ZAR1α1^), where the ZAR1 α1 helix was substituted for the NRC N-terminal equivalent sequence. We aligned the NRC2, NRC3, and ZAR1 α1 helix to determine where to make the chimaera junction (**Figure 4A**). To test whether these chimaeras can complement Cf-4/Avr4 or Pto/AvrPto-triggered cell death, we transiently expressed these receptor/effector pairs with either wild type NRC2 or NRC3, or their respective ZAR1 α1 helix chimaeras in *N. benthamiana nrc2/3/4* lines (**Figure 4B**). The chimeric proteins expressed and accumulated to similar level as both wildtype NRC2 and NRC3 proteins (**Figure 4C**). Like wild type NRC3, NRC3^ZAR1α1^ complemented both Cf-4/Avr4 or Pto/AvrPto-triggered cell death in the *N. benthamiana nrc2/3/4* CRISPR lines (**Figure 4D, E**). Surprisingly, the NRC2^ZAR1α1^ chimaera complemented Cf-4/Avr4-triggered cell death to a similar extent as NRC3 (**Figure 4D, E**). The NRC2^ZAR1^ chimaera also complemented Pto/AvrPto-triggered cell death to a greater extent than wild type NRC2 (**Figure 4D, E**, **Figure S5A** for the statistical analysis). Importantly, this was not because the NRC ZAR1 α1 helix chimaeras are autoactive, as expressing them with either Cf-4 or Pto and an EV control did not result in cell death (**Figure 4D, E**). We conclude that the α1 helix of ZAR1, which is known to make a Ca^2+^-permeable membrane-permeable pore, can replace the NRC2 and NRC3 α1 helix for hypersensitive cell death signaling.

**Figure 4:**
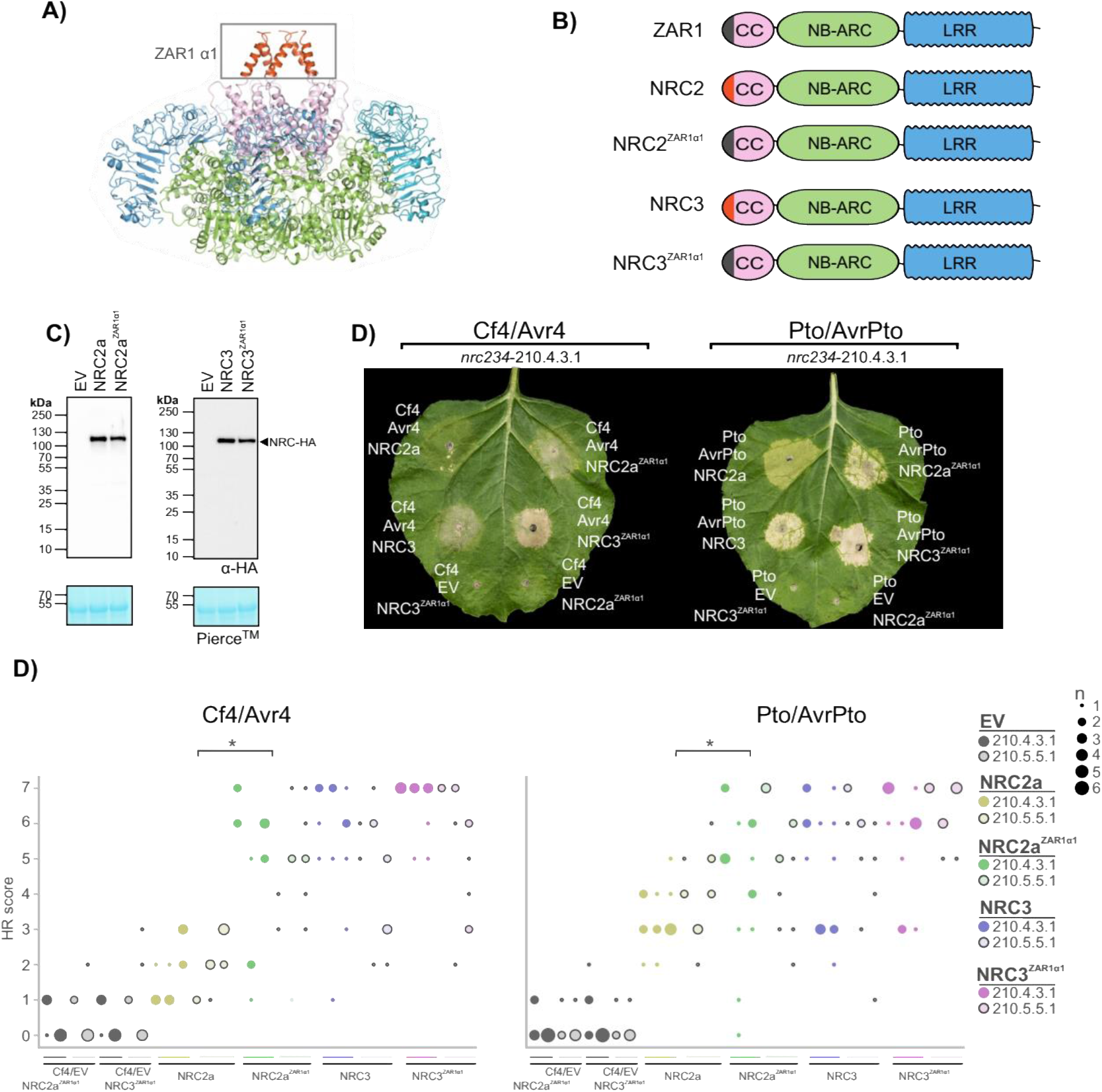
The ZAR1 α1 helix can functionally replace the NRC2 and NRC3 α1 helix for Cf-4/Avr4-triggered hypersensitive cell death. **A**) Structure of the ZAR1 resistosome with the N-terminal α1 helix highlighted. **B**) Schematic representation of ZAR1, NRC2, NRC3, and the respective NRC2^ZAR1α1^ and NRC3^ZAR1α1^ chimaeras. **C**) NRC ZAR1 α1 helix chimaeras accumulate to similar levels as wild type NRC proteins. Immunoblot analysis of transient NRC-6xHA accumulation 5 days after agroinfiltration in wild-type *N. benthamiana* plants. **D**) NRC ZAR1 α1 helix chimaeras can complement Cf-4/Avr4 and Pto/AvrPto-triggered hypersensitive cell death in the *N. benthamiana nrc234* CRISPR lines. Representative *N. benthamiana* leaves infiltrated with appropriate constructs were photographed 7 – 10 days after infiltration. The NRC tested, NRC2 and NRC3, are labelled above the leaf of NRC CRISPR line *nrc2/3/4*-210.4.3.1. The receptor/effector pair tested, Cf-4/Avr4 and Prf (Pto/AvrPto), are labelled on the leaf image. To ensure the NRC ZAR1 α1 helix chimaeras were not autoactive when expressed with either Cf-4 or Pto and an EV control. A representative leaf of the independent *nrc2/3/4*-210.5.5.1 CRISPR line is shown in **Figure S5**. **E**) Quantification of hypersensitive cell death. Cell death was scored based on a 0 - 7 scale between 7 – 10 days post infiltration. The results are presented as a dot plot, where the size of each dot is proportional to the count of the number of samples with the same score within each biological replicate. The experiment was independently repeated three times. The columns correspond to the different biological replicates. Significant differences between the conditions are indicated with an asterisk (*). Details of statistical analysis are presented in **Figure S5**.

### Pathogen effectors can suppress Cf4-triggered hypersensitive cell death in an NRC3-dependent manner

We recently showed that plant pathogens have converged on suppressing the NRC network to prevent immune responses (Derevnina *et al*., 2021). NRC3, for example, is suppressed by both the cyst nematode *Globodera rostochiensis* effector SPRYSEC15, as well as the oomycete *Phytophthora infestans* effector AVRcap1b (Derevnina *et al*., 2021). Since Cf-4/Avr4-triggered cell death requires NRC3, we reasoned that these effectors would also able to suppress cell death triggered by Cf-4 and similar LRR-RPs. To test this hypothesis, we generated heterologous expression constructs containing both NRC3 and either SPRYSEC15, AVRcap1b, or mCherry (control). We then co-expressed these NRC3/suppressor combinations together with Cf-4/Avr4 in *N. benthamiana nrc2/3/4* lines (**Figure 5A**). As controls, we used either autoactive NRC3^D480V^, or the receptor/effector pair Pto/AvrPto, both of which are suppressed by SPRYSEC15 and AVRcap1b (Derevnina et al., 2021). The receptor/effector pair R3a/Avr3a, which is not affected by SPRYSEC15 or AVRcap1b, was used as a control combination (Derevnina et al., 2021). These experiments revealed that both AVRcap1b and SPRYSEC15 suppress Cf-4/Avr4-triggered cell death (**Figure 5B, C**, **Figure S6A** for the statistical analysis). As expected, AVRcap1b and SPRYSEC15 suppressed autoactive NRC3^D480V^ and Pto/AvrPto-triggered cell death, but not R3a/Avr3a-triggered cell death (**Figure 5B, C**). As previously noted, transient expression of Cf-4/Avr4 results in a weak NRC3-independent chlorosis (**Figure 1**), and this phenotype was not suppressed by either SPRYSEC15 or AVRcap1b (**Figure 5B, C**). We conclude that divergent pathogen effectors, acting as suppressors of NRC3-mediated cell death, can suppress the hypersensitive cell death induced by the cell-surface receptor Cf-4.

**Figure 5:**
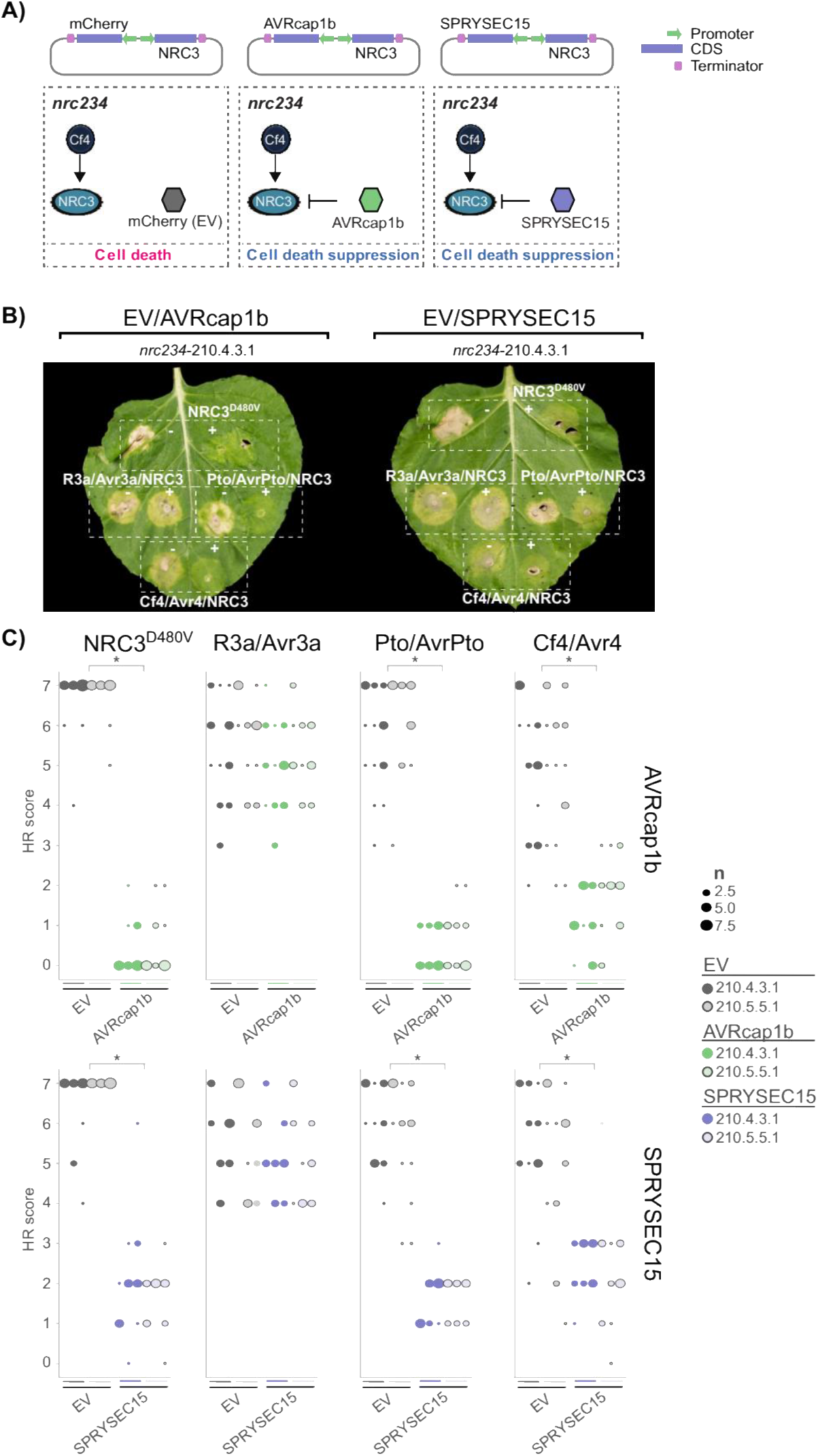
Cf-4/Avr4-triggered hypersensitive cell death can be suppressed by divergent pathogen effectors. **A**) A schematic representation of cell death assay strategy used. *A. tumefaciens* containing vectors carrying both NRC3 and either SPRYSEC15, AVRcap1b, or mCherry as empty vector control, were coexpressed in *N. benthamiana nrc2/3/4* CRISPR lines. Co-expression with NRC3-dependent receptor/effector pairs should result in suppressor-dependent suppression. **B**) Cf-4/Avr4-triggered hypersensitive cell death is suppressed by SPRYSEC15 and AVRcap1b, but not the EV control. Representative image of *N. benthamiana nrc2/3/4*-210.4.3.1 CRISPR line leaves which were agroinfiltrated with NRC3/suppressor constructs, as indicated above the leaf, and either autoactive NRC3^D480V^, Prf (Pto/AvrPto), R3a/Avr3a, or Cf-4/Avr4 as labelled on the leaf image, photographed 7 – 10 days after infiltration. A representative leaf of the independent *nrc2/3/4*-210.5.5.1 CRISPR line is shown in **Figure S6**. **C**) Quantification of hypersensitive cell death. Cell death was scored based on a 0 - 7 scale between 7 – 10 days post infiltration. The results are presented as a dot plot, where the size of each dot is proportional to the count of the number of samples with the same score within each biological replicate. The experiment was independently repeated three times. The columns correspond to the different biological replicates. Significant differences between the conditions are indicated with an asterisk (*). Details of statistical analysis are presented in **Figure S6**.

## DISCUSSION

Cell surface receptor-mediated immunity and intracellular NLR-mediated immunity are viewed as the main immune responses in plants. Although their signaling pathways were initially thought to be qualitatively distinct, recent findings point to a degree of cross-talk (Ngou *et al*., 2021; Yuan *et al*., 2021; Pruitt *et al*., 2021; Tian *et al*., 2021). Here we show that the cell-surface leucine-rich repeat receptor-like protein Cf-4 taps into the NRC helper NLR network to trigger hypersensitive cell death upon recognition of the apoplastic fungal *Cladosporium fulvum* effector Avr4. We propose that NRC3 is a core node that connects cell surface and intracellular immune networks (**Figure 6**). Divergent plant pathogens have convergently evolved to target the NRC3 node, thereby suppressing both intracellular and cell surface triggered immune responses (**Figure 6**).

**Figure 6:**
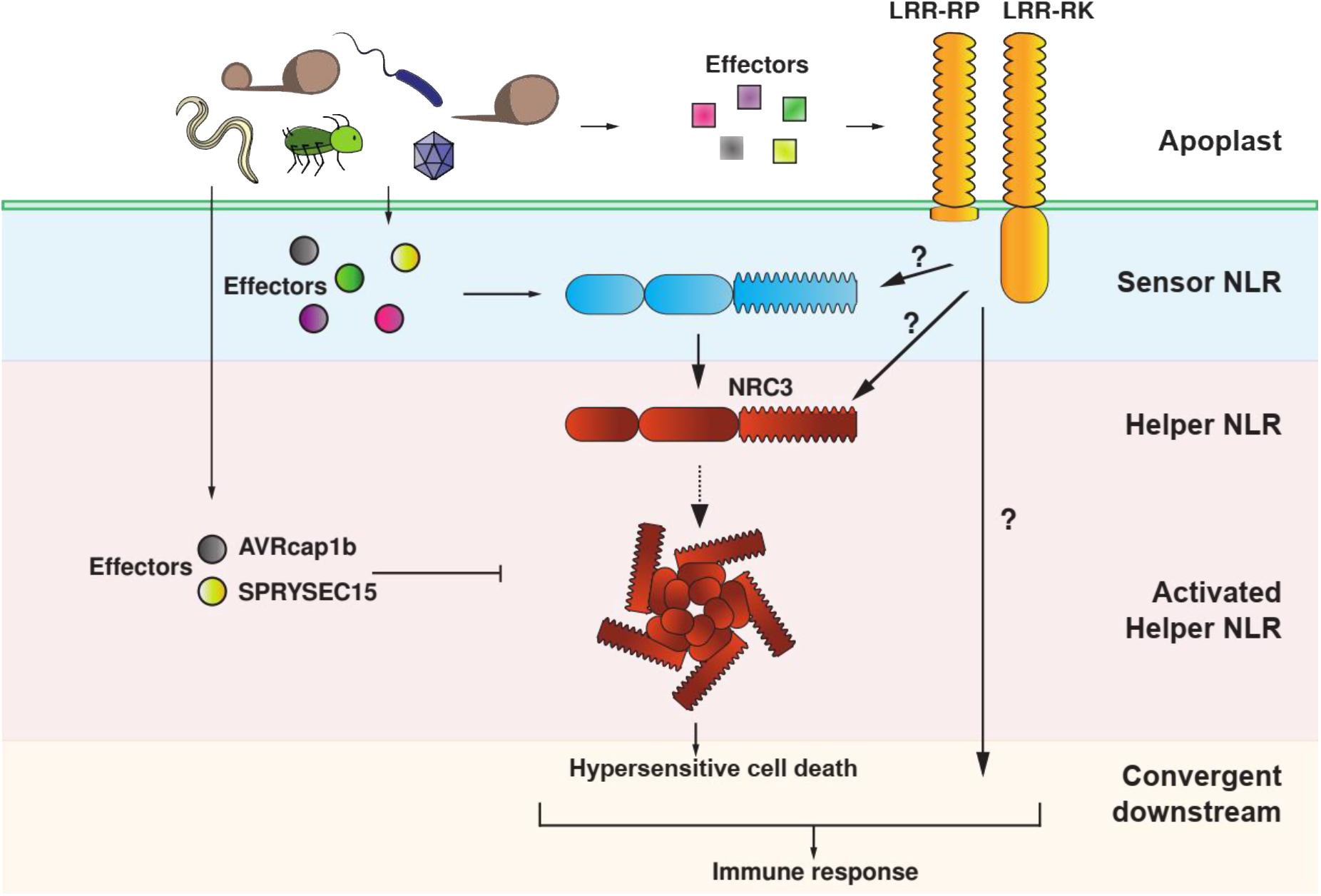
Hypersensitive cell death triggered by activation of cell-surface immune receptors requires the NRC network in Solanaceae. Recognition of apoplastic effectors and translocated intracellular effectors by cell-surface leucine-rich repeat receptor proteins and intracellular NRC-type sensor-NLRs, respectively, results in NRC3-mediated hypersensitive cell death. This cell death requires an intact α1 helix of NRC3 indicating the formation of an activated NRC3 resistosome. Divergent pathogen effectors converge on this node to suppress both NLR triggered as well as cell-surface receptor triggered immune recognition.

The hypersensitive cell death elicited by Cf-4/Avr4 is abolished in *N. benthamiana nrc2/3* and *nrc2/3/4*, but not *nrc4* lines (**Figure 1**). Interestingly, transient expression of Cf-4/Avr4 does trigger a NRC2/3/4-independent chlorosis (**Figure 1**) indicating that there is probably both NRC-dependent and -independent responses triggered by Cf-4, possibly mediated by the complex and dynamic association of RPs such as Cf-4 with partner RKs and RLCKs (Liebrand *et al*., 2013). Complementation using NRC3 restores Cf-4/Avr4-triggered hypersensitive cell death (**Figure 2**). NRC3 is conserved in all Solanaceous species investigated (**Figure S2**) and may mediate cell death responses triggered by cell-surface receptor in these species as well. When in evolutionary time this member of the NRC helper NLR family evolved to connect to the signaling pathway triggered by pattern recognition receptors remains to be elucidated.

Previously, NRC1 was implicated in the cell death mediated by the cell surface receptors Cf-4 (Gabriels *et al*., 2006), LeEIX2 (Gabriëls *et al*., 2007) and Ve1 (Fradin *et al*., 2009). Here we show that it is not NRC1, but NRC3 which links cell-surface immune receptors to intracellular immunity in the *N. benthamiana* experimental system (**Figure 1, 2**). Additionally, *Sl*NRC1 cannot complement Cf-4/Avr4-triggered hypersensitive cell death in the *N. benthamiana* experimental system (**Figure 2**). Unlike *Sl*NRC1, NRC3 is conserved in all Solanaceous species (**Figure S2**), indicating that this link between cell surface immune receptors and intracellular immunity may be conserved in the Solanaceae. In addition to the NRC-dependent hypersensitive cell death, there also appear to be NRC-independent responses downstream of these cell surface immune receptors. It may be that for some pathogens the NRC-independent response is sufficient to result in immunity, while for other pathogens the NRC-dependent response is also required. Which genetic components are involved in the NRC-independent response downstream of cell surface receptors remains a topic of further study.

NRC3 requires an intact MADA motif to mediate Cf-4/Avr4-triggered hypersensitive cell death and the ZAR1 α1 helix can substitute for the NRC3 α1 helix (**Figure 3**). This is similar to intracellular activation of NRCs, where NRC4-mediated Rpi-blb2/AVRblb2-triggered cell death and immunity requires an intact NRC4 MADA motif (Adachi, Contreras, *et al*., 2019). Additionally, the ZAR1 α1 helix can functionally substitute for the N-terminal equivalent sequence of NRC3, This observation suggests that the hypersensitive cell death response downstream of Cf-4/Avr4 immune signaling probably involves an activated resistosome.

Surprisingly, swapping the ZAR1 α1 helix into NRC2 resulted in a generally stronger cell death response as compared to the native NRC2. In contrast to the native NRC2, this chimeric protein could efficiently complement Cf-4/Avr4-triggered hypersensitive cell death (**Figure 4**). Given that the stronger immune responses triggered by the NRC2^ZAR1α1^ chimaera are not due to enhanced protein accumulation, the specific mechanisms that explain these differences remain unclear (**Figure 4**). It may be that this chimeric protein is more “trigger-happy”, either due to reduced autoinhibition or enhanced capacity to generate the resistosome funnel. Often trigger-happy mutations give increased autoimmune responses (Segretin *et al*., 2014; Chapman *et al*., 2014; Giannakopoulou *et al*., 2015). However, we did not observe autoimmune responses with the NRC2^ZAR1α1^ chimaera. Alternatively, the α1 helix may result in a more stable activated resistosome or form a more effective Ca^2+^ channel than the NRC3 N-terminus (see below). It may also be involved in subcellular localization and targeting either prior to, or after activation, and this localization could affect the strength of the immune response. Regardless of the mechanism, this chimeric approach has biotechnological implications and may be useful in engineering trigger-happy helper NLRs, such as the NRCs, that can mediate more potent immune responses.

The recent observation that the ZAR1 α1 helix forms a Ca^2+^-permeable membrane-permeable pore (Bi *et al*., 2021), and that the ZAR1 α1 helix can functionally substitute for the NRC2/3 N terminus, indicates that a Ca^2+^-permeable membrane-permeable pore is possibly involved in mediating Cf-4/Avr4-triggered cell death. It may be that this signaling is either a cell autonomous response directly downstream of activated cell surface receptors, or a response involved in amplification of the initial signaling response in either a cell autonomous or non-autonomous fashion. Genetic dissection of the NRC-independent responses may yield answers to these questions. It could very well be that upon activation of PRRs by pathogen molecules, multiple Ca^2+^ channels are activated during various stages in the signaling cascade, thereby explaining previous difficulties in genetically dissecting this response (Tian *et al*., 2019; Thor *et al*., 2020; Bjornson *et al*., 2021; Pruitt *et al*., 2021; Jacob *et al*., 2021).

We show that two highly divergent pathogen effectors which we recently characterized as suppressors of NRC3-mediated immune responses (Derevnina *et al*., 2021) can also compromise cell surface immune signaling (**Figure 6**). Therefore, in addition to suppressing intracellular immune recognition, SPRYSEC15 and AVRcap1b can also suppress immune recognition triggered by cell-surface receptors. Perhaps this is a more common approach by pathogens. We hypothesize that various cell surface receptors that are involved in nematode and oomycete immunity would also be dependent on helper NLRs of the NRC family similar to the fungal resistance protein Cf-4.

We have previously shown that the leucine-rich repeat receptor-like kinase FLS2 does not require NRC2/3/4 for triggering MAPK phosphorylation and a ROS burst upon recognition of the bacterial flagellin epitope flg22 (Wu *et al*., 2020). Furthermore, Suppressor of the G2 allele of skp1 (SGT1)-a core component of the CC-NLR-mediated hypersensitive cell death-is not required for receptor-like kinase signaling in *N. benthamiana* or Arabidopsis (Yu *et al*., 2021), further confirming that NRCs are not always downstream of leucine-rich repeat receptor kinases. It therefore seems likely that NRC3 is only required for full activation of the subset of cell surface receptors that can trigger hypersensitive cell death. Since NRCs are transcriptionally upregulated upon activation of Cf-4, FLS2 and other cell surface immune receptors (Gabriëls *et al*., 2006) it may be that Cf-4 activation results in transcriptional upregulation of NRC3 in an NRC-independent manner, to amplify the immune response and trigger hypersensitive cell death in a second phase in a NRC3-dependent manner.

Intracellular NLR immune receptors were recently shown to require intact cell surface immune signaling in order to efficiently trigger hypersensitive cell death (Yuan *et al*., 2021; Ngou *et al*., 2021). However, the mechanistic links between cell surface immune signaling and the activation of intracellular immune receptors remain unclear (Bjornson and Zipfel, 2021). Here we show that intracellular NLR immune receptors are also required for hypersensitive cell death induction by a cell surface receptor. In addition, we show that this hypersensitive cell death likely requires activation of a MADA-CC-NLR type resistosome. Understanding how these signaling modules intertwine and have evolved and how pathogens suppress these responses can be leveraged to guide new approaches for breeding disease resistance.

## MATERIAL & METHODS

### Plant material and growth conditions

Wild type and mutant *N. benthamiana* were propagated in a glasshouse and, for most experiments, were grown in a controlled growth chamber with temperature 22–25°C, humidity 45–65% and 16/8 hr light/dark cycle. The *NRC* CRISPR lines used have been previously described: *nrc2/3*-209.1.3.1 and *nrc2/3*-209.3.3.1 (Witek *et al*., 2021), *nrc2/3/4*-210.4.3.1 and *nrc2/3/4*-210.5.5.1 (Wu *et al*., 2020), and *nrc4*-185.1.2.1 (Adachi, Contreras, *et al*., 2019).

### Plasmid constructions

The Golden Gate Modular Cloning (MoClo) kit (Weber *et al*., 2011) and the MoClo plant parts kit (Engler *et al*., 2014) were used for cloning, and all vectors are from this kit unless specified otherwise. Effectors, receptors and NRCs were cloned into the binary vector pJK268c, which contains the Tomato bushy stunt virus silencing inhibitor p19 in the backbone (Kourelis *et al*., 2020). Cloning design and sequence analysis were done using Geneious Prime (v2021.2.2; https://www.geneious.com). Plasmid construction is described in **Table S1**.

### Transient gene-expression and cell death assays

Transient gene expression in *N. benthamiana* were performed by agroinfiltration according to methods described by van der Hoorn *et al*. (2000). Briefly, *A. tumefaciens* strain GV3101 pMP90 carrying binary vectors were inoculated from glycerol stock in LB supplemented with appropriate antibiotics and grown O/N at 28 °C until saturation. Cells were harvested by centrifugation at 2000 × *g*, RT for 5 min. Cells were resuspended in infiltration buffer (10 mM MgCl2, 10 mM MES-KOH pH 5.6, 200 μM acetosyringone) to the appropriate OD600 (see **Table S1**) in the stated combinations and left to incubate in the dark for 2h at RT prior to infiltration into 5-week-old *N. benthamiana* leaves. HR cell death phenotypes were scored in a range from 0 (no visible necrosis) to 7 (fully confluent necrosis) according to Adachi *et al*. (2019) (**Figure S1a**).

### Protein immunoblotting

Six *N. benthamiana* leaf discs (8 mm diameter) taken 5 days post agroinfiltration were homogenised in extraction buffer [10% glycerol, 25 mM Tris-HCl, pH 7.5, 1 mM EDTA, 150 mM NaCl, 1% (w/v) PVPP, 10 mM DTT, 1x protease inhibitor cocktail (SIGMA), 0.2% IGEPAL^®^ CA-630 (SIGMA)]. The supernatant obtained after centrifugation at 12,000 x g for 10 min 4 °C was used for SDS-PAGE. Immunoblotting was performed with rat monoclonal anti-HA antibody (3F10, Roche) or mouse monoclonal anti-GFP antibody conjugated to HRP (B-2, Santa Cruz Biotech) in a 1:5000 dilution. Equal loading was validated by staining the PVDF membranes with Pierce Reversible Protein Stain Kit (#24585, Thermo Fisher).

### Bioinformatic and phylogenetic analyses of the NRC-helper family

NRC sequences were retrieved by BLASTP against the NCBI RefSeq proteomes using the tomato and *N. benthamiana* NRC proteins (*Sl*NRC1, *Sl*NRC2, *Sl*NRC3, *Sl*NRC4a, *Sl*NRC4b, *Nb*NRC2a, *Nb*NRC2b, *Nb*NRC3, *Nb*NRC4a, *Nb*NRC4b, *Nb*NRC4c) as a query. A single representative splice-variant was selected per gene. Full-length amino acid sequences were aligned using MAFFT (v7.450; Katoh *et al*., 2002) (**Supplemental data 1**). FastTree (v2.1.11; Price *et al*., 2010) was used to produce a phylogeny of the NRCs which were rooted on XP_00428175 (**Supplemental data 2**). The NRC phylogeny was edited using the iTOL suite (6.3; Letunic and Bork, 2019).

## ACKNOWLEDGEMENTS

We thank Aleksandra Białas, Ana Cristina Barragan Lopez, Clémence Marchal, Daniel Lüdke, Paul Crosnier, and Thorsten Langner for their helpful comments on the figures. We thank Hsuan Pai for drawing the *N. benthamiana* plant in figure 2, Daniel Lüdke for helpful comments on the manuscript, and Phil Robinson for photography.

## AUTHOR CONTRIBUTIONS

Conceptualization: JK, HA, CHW, SK; Data curation: JK, MPC, CHW; Formal analysis: JK; Investigation: JK, AH, MPC, CHW; Supervision: SK; Funding acquisition: SK; Project administration: SK; Writing initial draft: JK, SK; Editing: JK, MPC, AH, HA, LD, SK, CHW.

---

**Figure S1:**
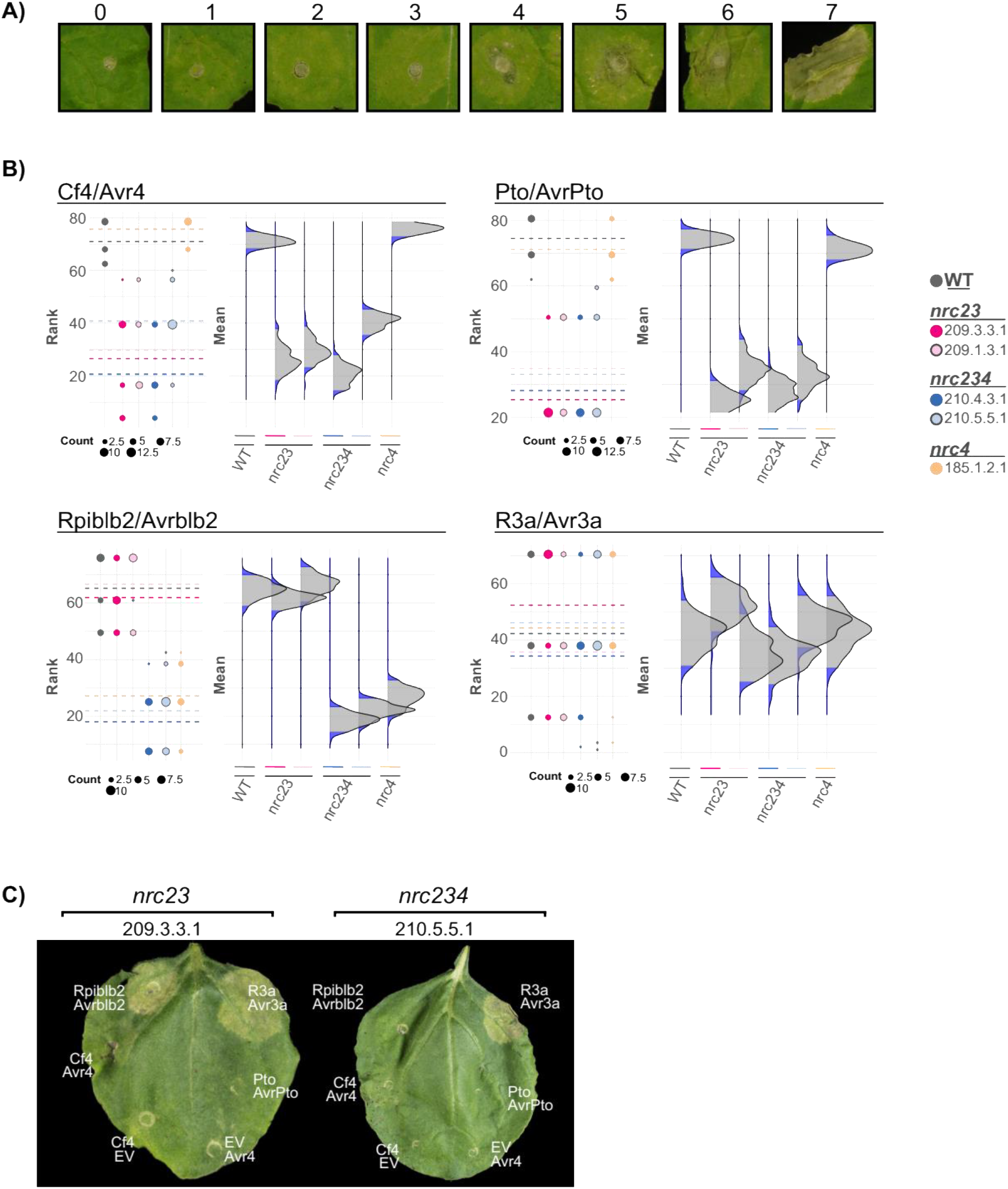
Statistical analysis of Cf-4/Avr4-triggered cell death in different *N. benthamiana nrc* mutant lines. **A)** Cf-4/Avr4-triggered cell death was scored on a scale of 0 – 7, with 0 being no response, and 7 being fully confluent cell-death in the entire infiltrated sector. Visible cell death starts appearing at a score of 4. **B)** Statistical analysis was conducted using the besthr R package. The dots represent the ranked data and their corresponding means (dashed lines), with the size of each dot proportional to the number of observations for each specific value (count key below each panel). The panels on the right show the distribution of 100 bootstrap sample rank means, where the blue areas under the curve illustrate the 0.025 and 0.975 percentiles of the distribution. A difference is considered significant if the ranked mean for a given condition falls within or beyond the blue percentile of the mean distribution of the wild-type control. **C)** Cf-4/Avr4-triggered hypersensitive cell death is lost in the *nrc2/3* and *nrc2/3/4* CRISPR lines. Representative *N. benthamiana* leaves infiltrated with appropriate constructs were photographed 7 – 10 days after infiltration. The NRC CRISPR lines, *nrc2/3*-209.3.3.1, *nrc2/3/4*-210.5.5.1, are labelled above the leaf and the receptor/effector pair tested, Cf-4/Avr4, Prf (Pto/AvrPto), Rpi-blb2/AVRblb2 or R3a/Avr3a, are labelled on the leaf image. Cf-4/EV and EV/Avr4 were also included.

**Figure S2:**
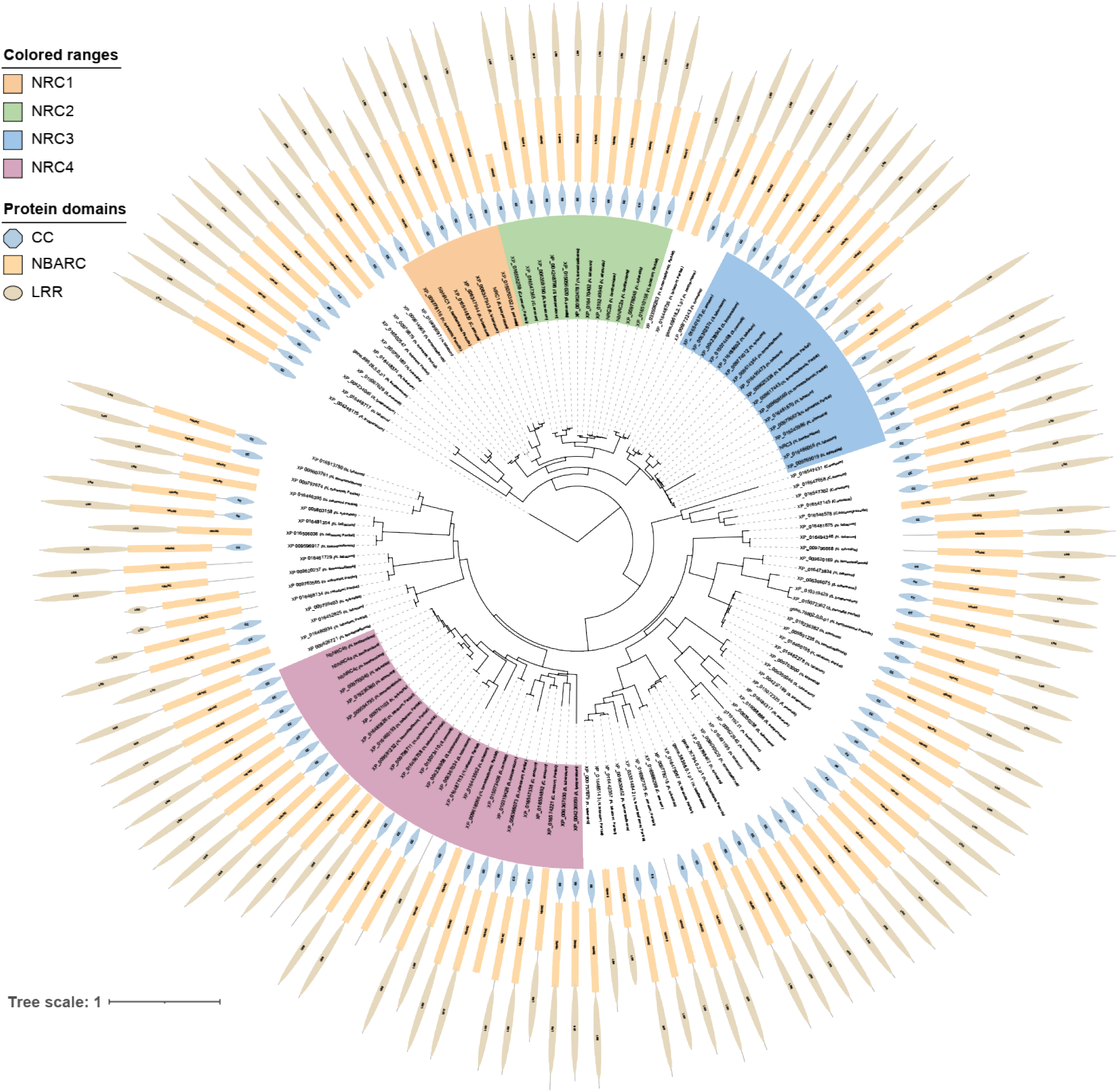
In contrast to the widely conserved NRC3, NRC1 is not present in *N. benthamiana*. Phylogenetic tree of the NRC-helper NLR family based on the full-length protein sequences was inferred using an approximately Maximum Likelihood method as implemented in FastTree (Price *et al*., 2010) based on the Jones-Taylor-Thornton (JTT) model (Jones *et al*., 1992). The tree was rooted on XP_00428175. The different NRC subfamilies are indicated. Domain architecture was determined using NLRtracker (Kourelis *et al*., 2021). Probable pseudogenes or partial genes, including *Nicotiana* NRC1 are indicated.

**Figure S3:**
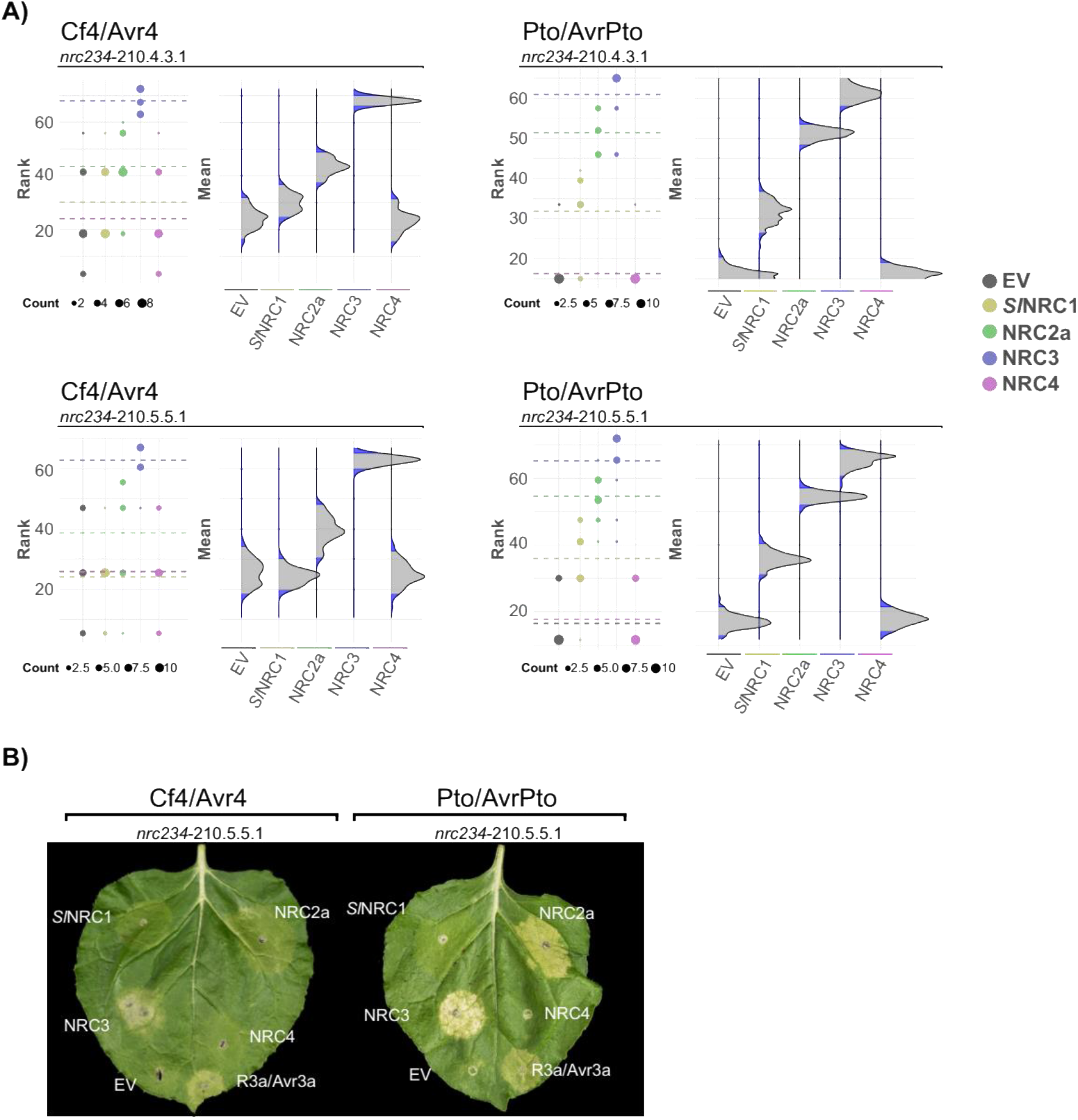
Statistical analysis of NRC complementation of Cf-4/Avr4-triggered cell death *N. benthamiana nrc2/3/4* CRISPR lines. **A**) Statistical analysis was conducted using besthr R package (MacLean, 2020). The dots represent the ranked data and their corresponding means (dashed lines), with the size of each dot proportional to the number of observations for each specific value (count key below each panel). The panels on the right show the distribution of 100 bootstrap sample rank means, where the blue areas under the curve illustrate the 0.025 and 0.975 percentiles of the distribution. A difference is considered significant if the ranked mean for a given condition falls within or beyond the blue percentile of the mean distribution of the wild-type control. **B**) Cf-4/Avr4-triggered hypersensitive cell death is complemented in the *nrc2/3/4* CRISPR lines with NRC3 and to a lesser extent NRC2, but not NRC1 or NRC4. Representative *N. benthamiana* leaves infiltrated with appropriate constructs were photographed 7 – 10 days after infiltration. receptor/effector pair tested, Cf-4/Avr4 and Prf (Pto/AvrPto), are labelled above the leaf of NRC CRISPR line *nrc2/3/4*-210.5.5.1. The NRC used for complementation or EV control are labelled on the leaf image.

**Figure S4:**
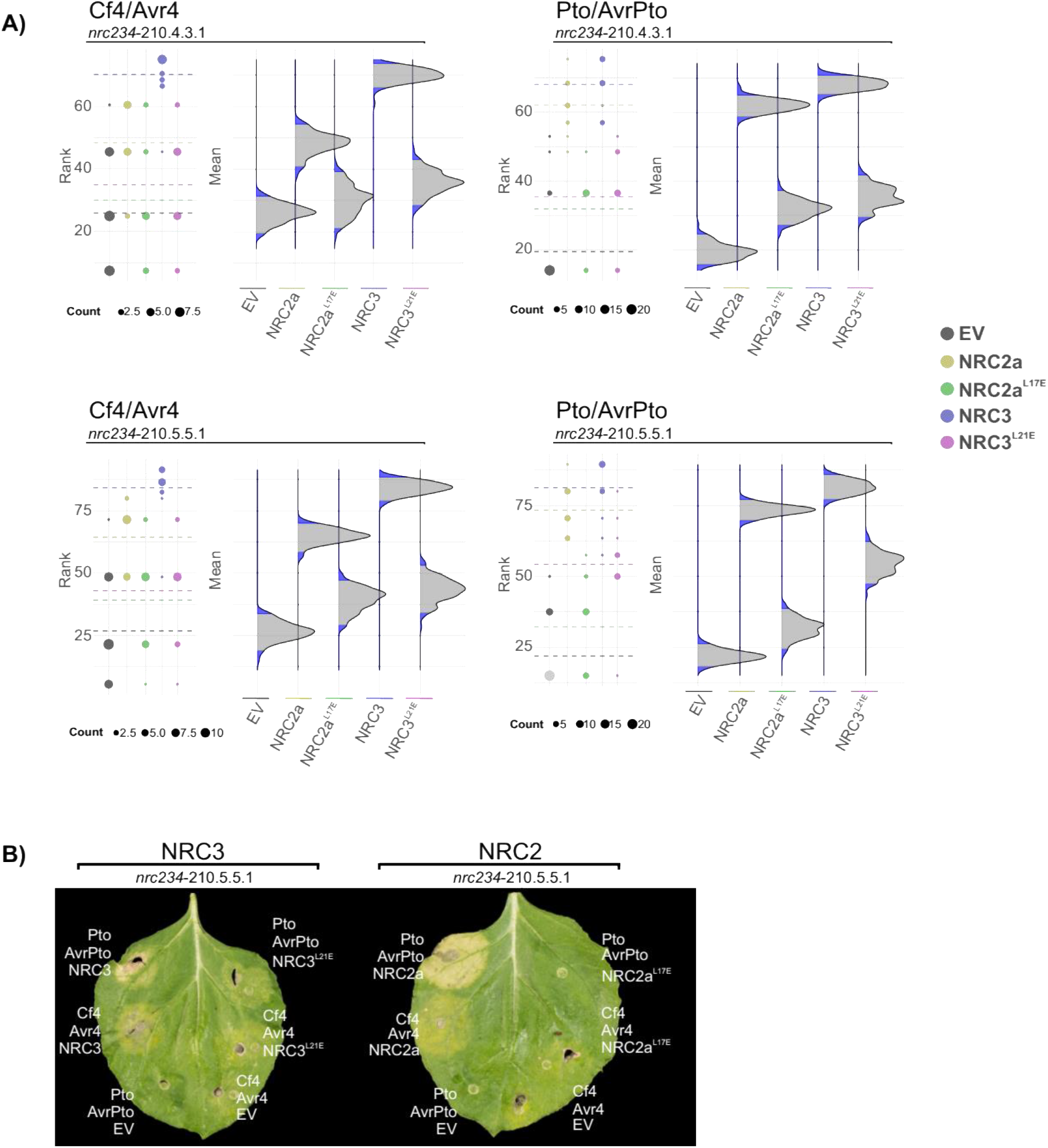
Statistical analysis of NRC MADA mutant complementation of Cf-4/Avr4-triggered cell death *N. benthamiana nrc2/3/4* CRISPR lines. **A**) Statistical analysis was conducted using besthr R package (MacLean, 2020). The dots represent the ranked data and their corresponding means (dashed lines), with the size of each dot proportional to the number of observations for each specific value (count key below each panel). The panels on the right show the distribution of 100 bootstrap sample rank means, where the blue areas under the curve illustrate the 0.025 and 0.975 percentiles of the distribution. A difference is considered significant if the ranked mean for a given condition falls within or beyond the blue percentile of the mean distribution of the wild-type control. **B**) NRC MADA mutants cannot complement Cf-4/Avr4 or Pto/AvrPto-triggered hypersensitive cell death in the *N. benthamiana nrc2/3/4* CRISPR lines. Representative *N. benthamiana* leaves infiltrated with appropriate constructs were photographed 7 – 10 days after infiltration. The NRC tested, NRC2 and NRC3, are labelled above the leaf of NRC CRISPR line *nrc2/3/4*-210.5.5.1. The receptor/effector pair tested, Cf-4/Avr4 and Prf (Pto/AvrPto), are labelled on the leaf image. To ensure the NRC MADA mutants were not autoactive they were expressed with either Cf-4 or Pto and an EV control.

**Figure S5:**
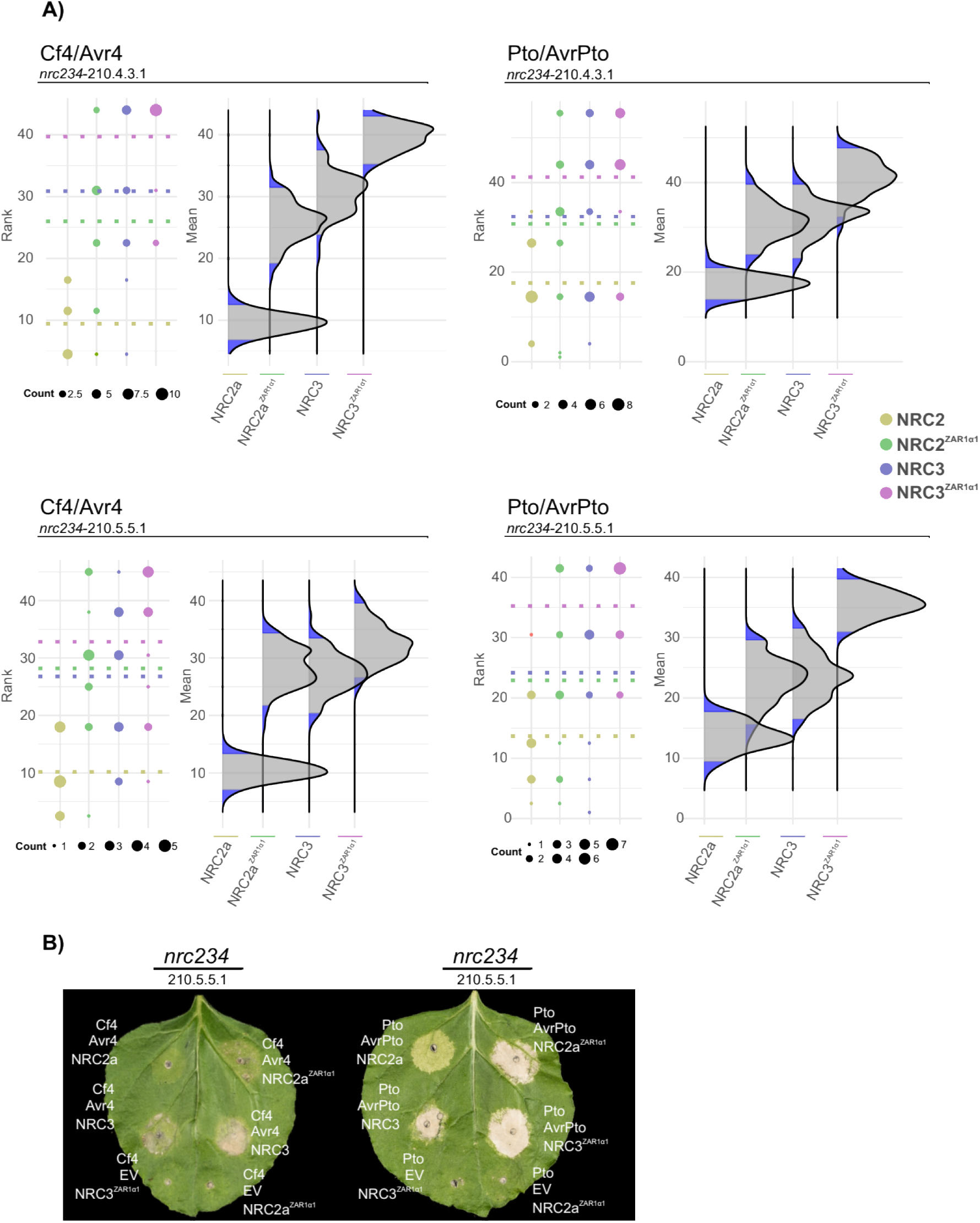
Statistical analysis of NRC ZAR1 α1 helix chimaera complementation of Cf-4/Avr4-triggered cell death *N. benthamiana nrc2/3/4* CRISPR lines. **A**) Statistical analysis was conducted using besthr R package (MacLean, 2020). The dots represent the ranked data and their corresponding means (dashed lines), with the size of each dot proportional to the number of observations for each specific value (count key below each panel). The panels on the right show the distribution of 100 bootstrap sample rank means, where the blue areas under the curve illustrate the 0.025 and 0.975 percentiles of the distribution. A difference is considered significant if the ranked mean for a given condition falls within or beyond the blue percentile of the mean distribution of the wild-type control. **B**) NRC ZAR1 α1 helix chimaeras can complement Cf-4/Avr4 and Pto/AvrPto-triggered hypersensitive cell death in the *N. benthamiana nrc2/3/4* CRISPR lines. Representative *N. benthamiana* leaves infiltrated with appropriate constructs were photographed 7 – 10 days after infiltration. The NRC tested, NRC2 and NRC3, are labelled above the leaf of NRC CRISPR line *nrc2/3/4*-210.5.5.1. The receptor/effector pair tested, Cf-4/Avr4 and Prf (Pto/AvrPto), are labelled on the leaf image. To ensure the NRC ZAR1 α1 helix chimaeras were not autoactive they were expressed with either Cf-4 or Pto and an EV control.

**Figure S6:**
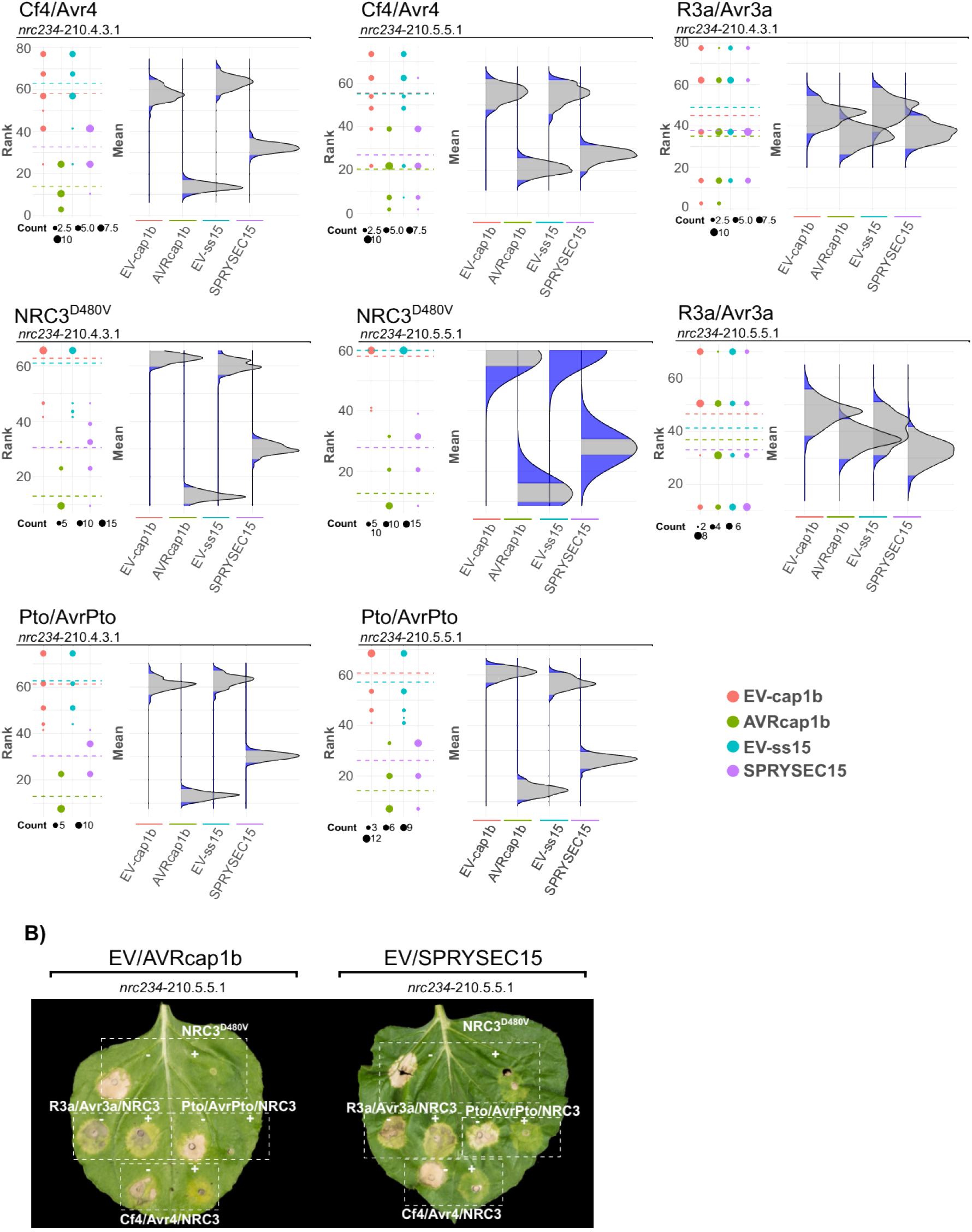
Statistical analysis of Cf-4/Avr4-suppression mediated by divergent pathogen effectors. **A**) Statistical analysis was conducted using besthr R library (MacLean, 2020). The dots represent the ranked data and their corresponding means (dashed lines), with the size of each dot proportional to the number of observations for each specific value (count key below each panel). The panels on the right show the distribution of 100 bootstrap sample rank means, where the blue areas under the curve illustrates the 0.025 and 0.975 percentiles of the distribution. A difference is considered significant if the ranked mean for a given condition falls within or beyond the blue percentile of the mean distribution of the wild-type control. **B**) Cf-4/Avr4-triggered hypersensitive cell death is suppressed by SPRYSEC15 and AVRcap1b, but not the EV control. Representative image of *N. benthamiana nrc2/3/4*-210.5.5.1 CRISPR line leaves which were agroinfiltrated with NRC3/suppressor constructs, as indicated above the leaf, and either autoactive NRC3^D480V^, Prf (Pto/AvrPto), R3a/Avr3a, or Cf-4/Avr4 as labelled on the leaf image, photographed 7 – 10 days after infiltration.

## SUPPLEMENTAL DATA

**Table S1: Description of constructs and *Agrobacterium* strains.**

**Supplemental dataset 1: MAFFT alignment of the unique full-length NCBI RefSeq NRCs (Fasta format).** This file contains the MAFFT alignment of 134 unique NRCs found in the NCBI RefSeq database. NRC0 was added as an outgroup.

**Supplemental dataset 2 Approximately maximum likelihood phylogenetic tree of the NCBI RefSeq NRCs (Nexus format).** This file contains the phylogenetic analysis of the full-length NRCs found in the NCBI RefSeq database using the JTT method.

